# Time to degree, identity, and grant writing: Lessons learned from a mixed-methods longitudinal case study

**DOI:** 10.1101/2023.01.26.525758

**Authors:** AJ Alvero, Courtney Peña, Amber R Moore, Leslie Luqueno, Cisco B. Barron, Latishya Steele, Stevie Eberle, Crystal M. Botham

## Abstract

Time to degree completion is an important metric of academic progress and success for doctoral students. It is also a common way for educational stakeholders to compare programs even if the content of the degree programs varies. But what types of behaviors and experiences are associated with faster times to degree? In this education article, we examine the relationship between receiving competitive research awards (e.g. grant writing) and time to degree for PhD students. We organize our analyses by student identities, specifically gender and underrepresented minority (URM) status, to examine differences in time to degree based on student demographics. Our dataset included students that graduated between academic years 2008-09 through 2018-19. We also interviewed students currently enrolled in these same programs. We find that URM women who won competitive research awards graduate faster than all other students who also won awards but also report lower levels of advisor support. We also find that URM women and all URM students tended to graduate faster compared to other groups of students who did not win awards. Students who reported feeling supported by their advisors, most of which reflected hands-on guidance through the writing process, in the interviews were more likely to apply for grants. Combined, these results highlight that writing grants and specific types of advisor support may influence faster times to degree for bioscience PhD students. This study suggests similar introspective analyses at other institutions and databases are needed.

## 1 Introduction

Writing is, in some ways, the most essential skill that students develop and fine tune during their time in PhD programs. Scientific communication is strongly linked to academic success including research career persistence (Cameron et al., 2020). Although most of the attention on academic writing is centered on publications and presentations, grant writing is important for doctoral students because it provides them an opportunity to demonstrate their ability to think and act like a scientist by developing an original research project (Files et al., 2020). And because a successful career in science, especially the biosciences, is so dependent on one’s ability to secure external funding, grants are a critical academic currency. Aside from the financial benefits of securing grants, grant writing builds skills like critical thinking and strategies for communicating scientific ideas (Kahn et al., 2016; Quitadamo & Kurtz, 2007). Grant writing is also essential for advancement at the faculty level and securing grants improves one’s chances of advancing to tenure (Bloch et al., 2014; Ransdell et al., 2021; Smith et al., 2017; Thorpe, et al., 2020). While the links between faculty career success and grant writing is evident, the relationship between PhD student success and grant writing is less understood. The ways that PhD advisors support their students and whether or not they correlate with positive grant writing outcomes are also not well documented.

There are many different markers for success as a PhD student, but degree completion is a significant one (de Valero, 2001). The time it takes to complete a degree can relate to delayed career earnings, use of institutional resources, and differing amounts of student debt (e.g. accrued interest on student loans). In this way, a student’s time to degree, measured by the number of active graduate quarters (or semesters) until PhD conferral, is a significant commitment for students as they develop professionally and is also important to the institutions that support them. It is also a metric used by doctorate granting universities to compare outcomes (Blank et al., 2017). Other studies have considered factors like marital status, race, gender, academic performance (undergrad GPA and GRE scores) on time to degree and other measures of academic accomplishment (Akabas & Brass, 2019; Feldon et al., 2017; Mendoza-Sanchez et al., 2022; Petersen et al., 2018; Price, 2006), but the relationship between advanced academic tasks like grant writing and time to degree is less understood. Therefore, if faculty success is strongly associated with winning grants, how might winning grants be associated with PhD student success in terms of time to degree? Some data indicates that submitting or receiving external funding impacts time to degree (Ehrenberg & Mavros, 1995) but no formal data has yet been collected or analyzed to support this specifically for the biosciences. To bridge this gap, this study assessed the impact of external funding on time to degree for doctoral students studying the biosciences at Stanford University and which types of advisor support helps students secure external funding once they are enrolled.

Doctoral experiences, like all social experiences, are shaped by one’s personal identity in ways that reflect patterns of inequity in STEM (Bernard & Cooperdock, 2018; Chaudhary & Berhe, 2020; Committee on Effective Mentoring in STEMM et al., 2019; Master & Meltzoff, 2020; McCoy et al., 2017; Palmer et al., 2011). Student diversity based on race and gender is important for inclusion and representation, but determining factors associated with their success is also important. In some cases, student diversity and identity have been sources of intellectual innovation despite these same students receiving less recognition (Hofstra et al., 2020). Other studies have observed that STEM students from minoritized backgrounds are also less likely to feel senses of belonging in ways that are associated with positive academic outcomes (Walton et al., 2012; Walton & Cohen, 2007, 2011) and negative mental health symptoms (Sargent et al., 2002; Strayhorn TL, 2018). Given the stakes and prior research, our paper weaves these different threads together using longitudinal and interview data to answer question about the relationship between securing external funding and time to degree while also tending to whether student identities are salient in these relationships. Our paper answers the following questions about the nature of the relationship between time to degree, student identity, and winning competitive grants and fellowships:

- What is the relationship between time to degree and winning competitive awards for students in the biosciences?
- What are the ways students feel supported by their advisors (or not) in the grant writing process? Are there specific sets of behaviors associated with feelings of support as well as success in the grant writing process?
- Does the level of advisor support vary along dimensions of student identity, specifically race and gender?

We find that winning competitive awards is associated with a shorter time to degree but only for specific groups of students, including URM women. We also find that URM women are less likely to feel supported by their advisors to pursue grant writing, and that non-URM students were more likely to feel supported and apply for external funding. These findings, which we describe in more detail below, show how an important aspect of professional success among faculty (e.g. grant writing) is also correlated with how long it takes for students to complete their PhD degrees.

## 2 Materials and Methods

Our data come from Stanford University bioscience PhD programs in two forms: longitudinal data from an internal database and from interview and survey data. The data include information about the student’s gender, specific program, citizenship status, and time to degree. See Table 1 for a breakdown of the longitudinal data. To ensure participant anonymity, we do not provide a full breakdown of the interview participants. To study the relationships between time to degree and funding acquisition, we considered the students’ gender and URM status. We defined URM using federal categories mandated by the Department of Education; this is also how large organizations like NIH define URM status. The URM category includes students who are US citizens or permanent residents and who self-identify as American Indian/Alaska Native, Black/African American, Hispanic/Latino, or Native Hawaiian/Other Pacific Islander. The category includes any student who affiliates, with any one of these categories, regardless of whether they also affiliate with other categories (i.e., affiliating as “White” and “Black/African American” yields a URM designation). These same criteria were used in the survey and interview data to determine who is URM. We excluded students who transferred into the program or discontinued before graduating (n = 12). Medical Scientist Training Program (MSTP) students were not included in our analyses. We used Stanford’s Student Integrated Reporting and Information System (SIRIS) to query official university enrollment records related to time-to-degree and student demographics.

**Table 1:**
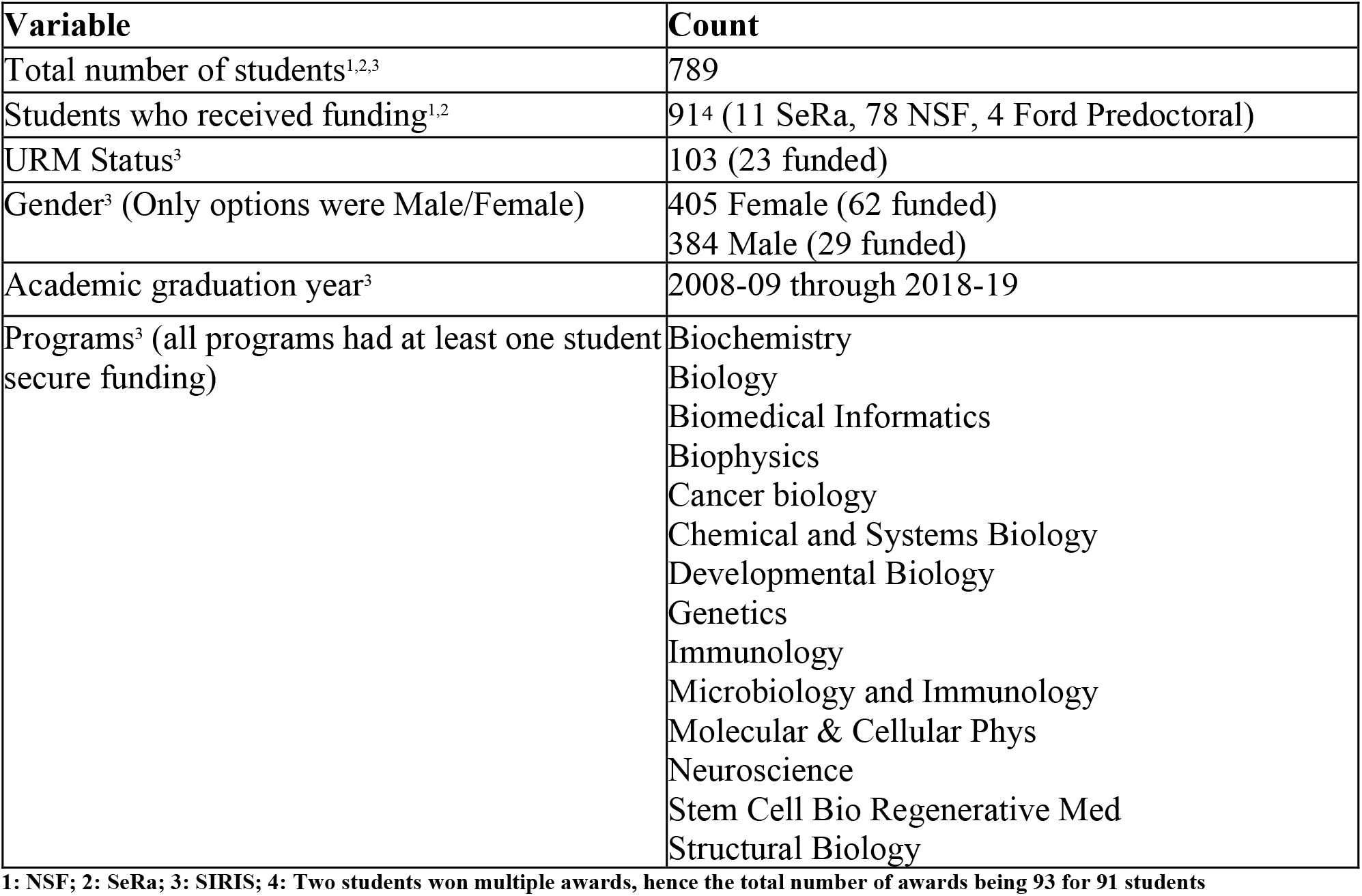
Overview of data.

Our quantitative data about student grant successes were pulled from several data sources, queried in September 2022. We leveraged the Stanford Electronic Research Administration System (SeRA), a database which tracks external proposals, such as fellowships or grants, submitted and awarded to grant seekers at Stanford (including doctoral students); see (Botham et al., 2020) for more information about the database. We also considered other fellowships that are not tracked by SeRA but are popular among students at Stanford (e.g. National Sciences Foundation (NSF) Graduate Research Fellowship, Ford Fellowship, Soros Fellowship, National Defense Sciences and Engineering Graduate (NDSEG) Fellowship). The publicly available NSF Graduate Research Fellowship Program (GRFP) Award Offers and Honorable Mentions List (see https://www.research.gov/grfp/) provided names of graduate students that won the NSF GRFP Award and Honorable Mentions status. The Ford Foundation Predoctoral Fellowship also posts the names of recipients (see Ford Fellows directory). Fellowships not based directly on research (Soros Fellowship) and with limited publicly available information (NDSEG Fellowship) were excluded. We wanted to provide this additional detail to encourage other researchers interested in replicating this study. Funding sources included the National Institutes of Health (NIH), the Howard Hughes Medical Institute, and more specialized sources such as the American Brain Tumor Association. We do not know how many of the students in our sample applied to the NSF GRFP, but 140 students applied for external funding and of those students 59 won awards (data not shown).

We use two statistical tests to compare average times to degree between different groups of students. First, we used Welch’s t-test to compare group means while accounting for different group sizes. Second, we used Cohen’s *d*, an effect size metric used often in education research (Allington et al., 2023; Kumari & Kumar, 2023). The outcome of interest that we compared is the elapsed time to degree measured in years. We grouped the students by URM status, gender, and the intersections of URM status and gender. Although we tested many different groups, we only present the results that were statistically significant with the t-testing (ie. p-value < 0.05). We indicated in figures and the text statistical significance with one asterisk indicating p-values below 0.05; two asterisks for p-values below 0.01; and three asterisks for p-values below 0.001. We also follow well established conventions when describing the effect sizes and present effect sizes that were not null (ie. 95% confidence interval does not include 0). To interpret the size of the effect, we refer to Cohen’s original suggestions where an effect size of 0.2 is small, 0.5 is medium, and .8 and above is large (Cohen, 2013). To prevent potential false discovery across the many comparisons within our data, we exclude any effect sizes that are below 0.5 (medium). We also filter out results that are not at least medium with respect to Cohen’s *d* and statistically significant according to the t-tests. Combining both allows for our analysis to go beyond p-values and think about effect sizes for more robust analysis and interpretation (Sullivan & Feinn, 2012). Part of our goal with this study is to encourage other institutions to look at their own data, and including the strongest results and comparisons that pass multiple tests of significance help us present more focused suggestions.

We draw qualitative data from a screener survey and interviews of doctoral students currently enrolled in the same bioscience programs. A recruitment email with a screener survey was sent to all bioscience PhD students who were enrolled in at least their third year. The screener survey was used to ensure diversity of interview participants and asked for basic demographic information, whether or not they applied for awards and whether or not they won. To maintain anonymity of participants, we do not specify which awards they applied for and/or won. After recruitment and screening, we completed 17 semi-structured interviews during the summer of 2022. From our interview pool, 76% applied for grants; 58% received grants; 24% were URM; and 18% were male. The qualitative data will allow us to answer our questions about factors associated with successful grant writing and which students are feeling supported.

The semi-structured interviews were transcribed and then analyzed using an inductive framework. The students were asked about their experiences in their respective programs, if and how they felt supported to pursue grant writing from their advisors, and their perceptions of the relationship between their personal identity and doctoral experiences. After transcribing the interviews, we did a thematic analysis (Braun & Clarke, 2012) to understand how student support could lead to positive grant writing experiences. Pairing the quantitative and qualitative analyses gives us a more holistic perspective on the relationship between grant writing and time to degree among bioscience PhD students.

Though we do not make any claims to generality or causality, we aim to encourage researchers at other institutions to investigate their own data and adapt a similar mixed methods approach to also capture the multiple perspectives we share here. Further, if a nationally representative dataset were to be created to explore factors affecting time to degree, such as grant writing, we would encourage investigation into such data using our approach but also standard tools for causal inference.

## 3 Results

### 3.1 Quantitative results: Longitudinal data

Among all students in the longitudinal dataset, the average time to degree was 5.86 years; the median time was 5.80 years; and the range was 3.0 to 11.5 years. See Figure 1 for a density plot of the years to degree. Although the range is wide, 68% of the PhD students in our data completed their programs in six or fewer years, similar to (though higher than) a national study of bioscience PhD students that reported 49% of students complete their degrees in 6 or fewer years (Ostriker et al., 2010). Next, we compared average times to degrees for different groups of students with respect to their identity. The first two analyses do not include the four Ford Predoctoral Fellowship winners in the “winner” groupings. The Ford Predoctoral winners had a higher average time to degree than other students (6.80 years) and, unlike the other funding programs considered, the program specifically targets underrepresented minorities in academia.

**Fig. 1.**
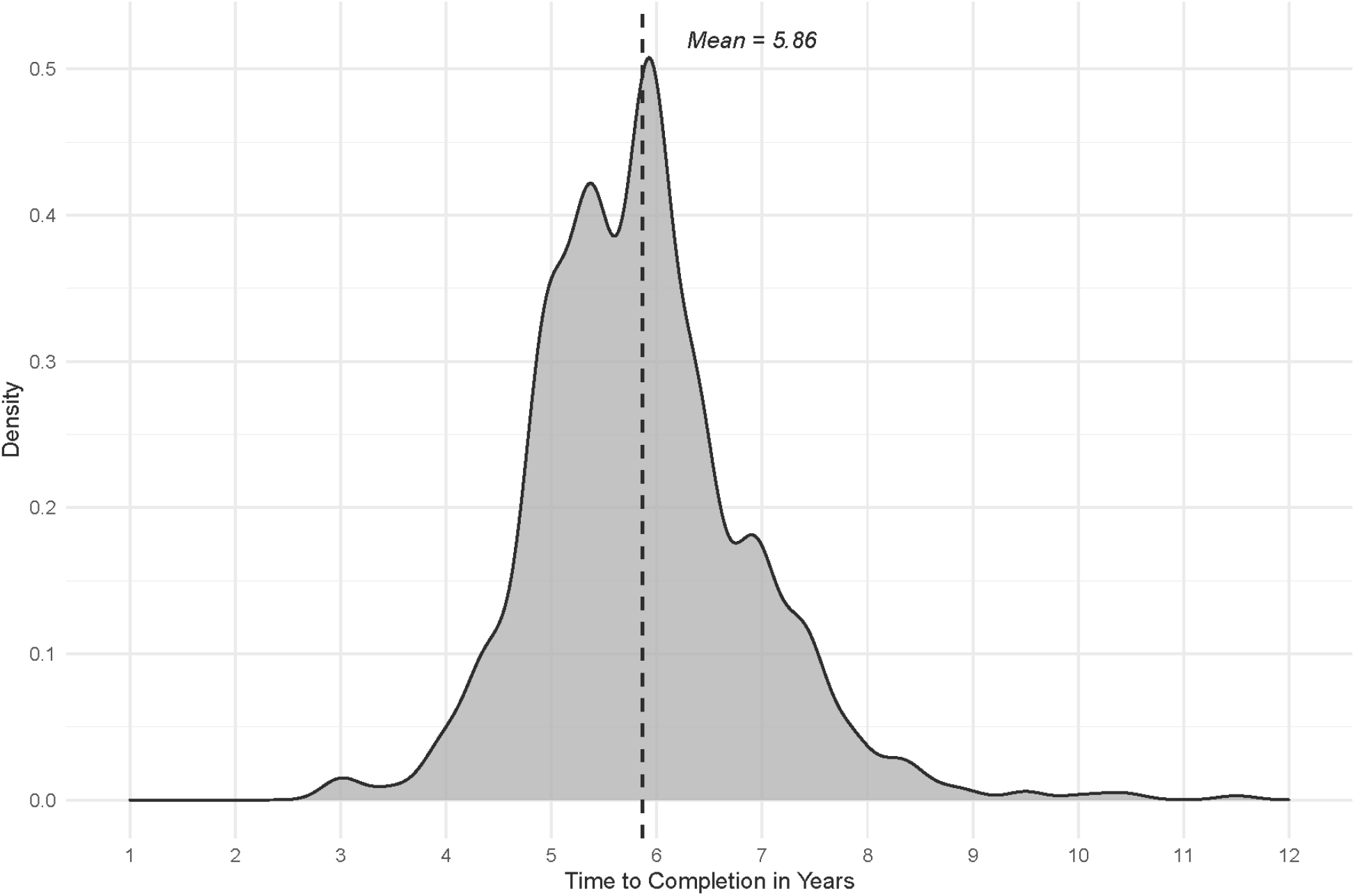
Density plot for years to degree, including the mean time to degree (5.86 years). The median time to degree (5.80 years) is not included to make the figure easier to interpret.

Our initial comparisons were based on student gender, using sex as a surrogate, or URM status. In our data, students could only select Male or Female. To be inclusive and accurately represent the distinction between sex and gender, future studies might consider non-binary students, but this is a limitation of our statistical analysis on student gender. None of the comparisons we tested based on student gender and winning grants were statistically significantly different. For example, there was no significant difference between women who won grants compared to women who did not (5.90 years and 5.84 years on average respectively; data not shown), nor was there a difference between men who won grants and women who won grants (5.71 years and 5.90 years on average respectively; data not shown).

This was not the case for student URM status, however. URM students who won grants graduate on average 0.53 years faster than non-URM students who also won grants (Figure 2A). And although gender in isolation was not strongly associated with faster or slower degree completion times, the intersection of URM status and gender did show a strong association. URM women who won grants graduate significantly faster than URM women who did not receive grants (5.15 years and 6.00 years median respectively, p value = 0.00008). Figure 2B shows a boxplot in which the fastest quartile of URM women who did not win grants graduated at around the same time as the slowest quartile among URM women who won grants. This result is also noteworthy when compared to average time to degree (5.86 years; Figure 1) for all the students in the dataset.

**Fig. 2a.**
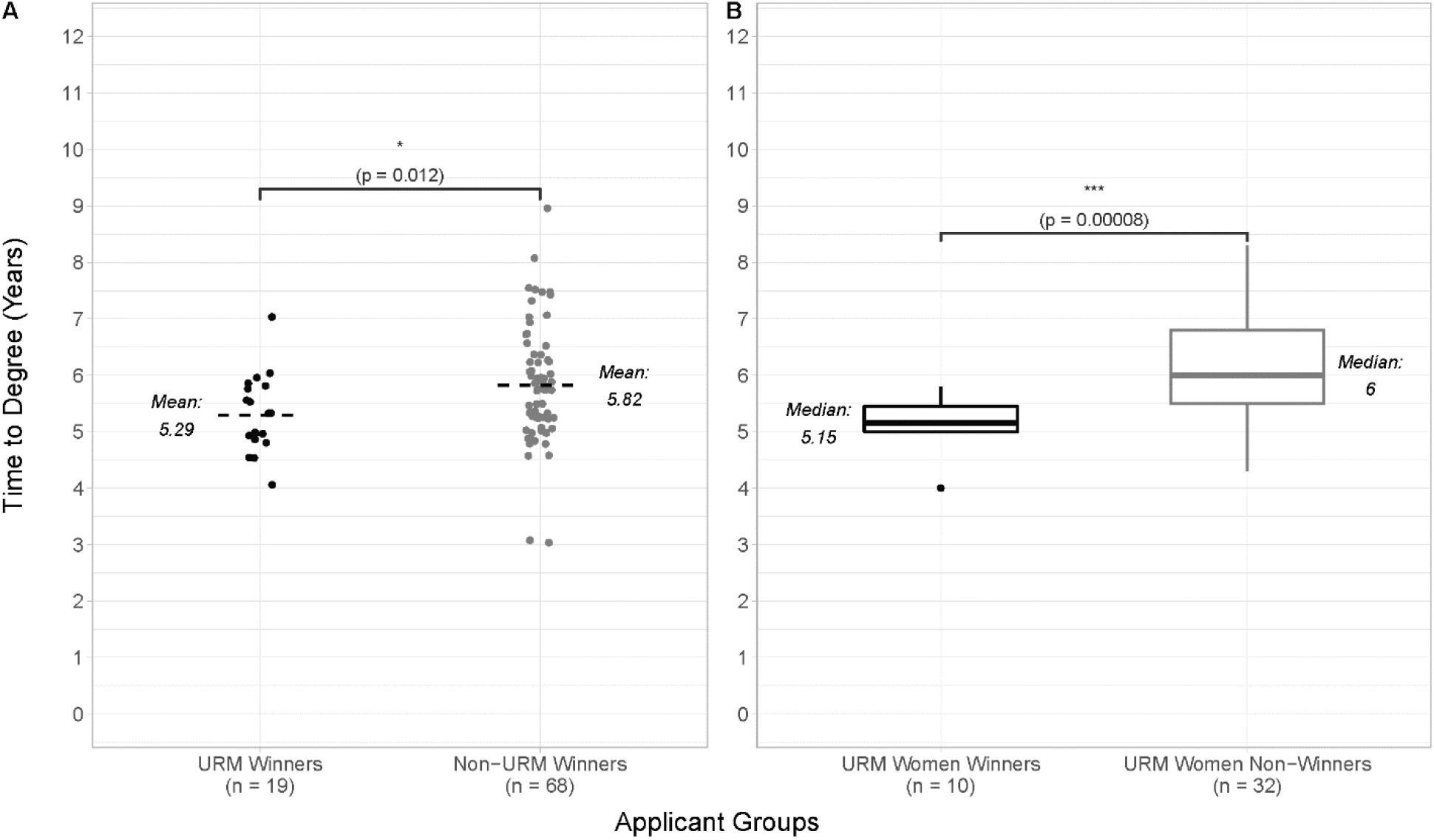
URM students who won grants (black dots) graduated faster than non-URM students who won grants (gray dots). The mean time to degree for each group is indicated by the dashed lines and the p-value is indicated by the asterisk at the top of the figure (p < 0.05). **Fig. 2b** Boxplot showing that URM women who win grants (black) graduate nearly one year faster on average than URM women who do not win grants (gray). The p-value is indicated by the asterisk at the top of the figure (p < 0.001).

We find URM winners graduated a full academic year (0.74 calendar years or about 9 months) faster on average than URM non-winners (Figure 3). The time to degree for the Ford Fellows ran counter to the other trends we found by taking an average of 6.80 years to complete their PhD. This was also on average 0.77 years (a full academic year) longer than URM non-winners and 1.51 years (more than two academic years) longer than URM winners (Figure 3). In our data set, the four students who won the Ford Fellowship did not also win NSF or SeRA-tracked funding. We included them within the URM non-winner group (see dark gray triangles) given the unique expectations for the Ford Fellowship on “sustained personal engagement with communities that are underrepresented in the academy” *Ford Foundation Predoctoral Fellowship Fact Sheet* (2023). Results show a strong relationship with time to degree and winning competitive awards but in the opposite direction for the Ford Fellowship: longer as opposed to shorter average times to degree. A significant aspect to participation in the Ford programs is service and volunteering, and the longer time to degree for these students might reflect the realities of trying to balance the work of advancing in a doctoral program with competing service goals. Although the Ford Foundation is ending their doctoral fellowship programs, this finding highlights that the relation between time to degree and winning competitive awards might not be purely transactional but also dependent on the nature and expectations that come attached to the funding.

**Fig. 3.**
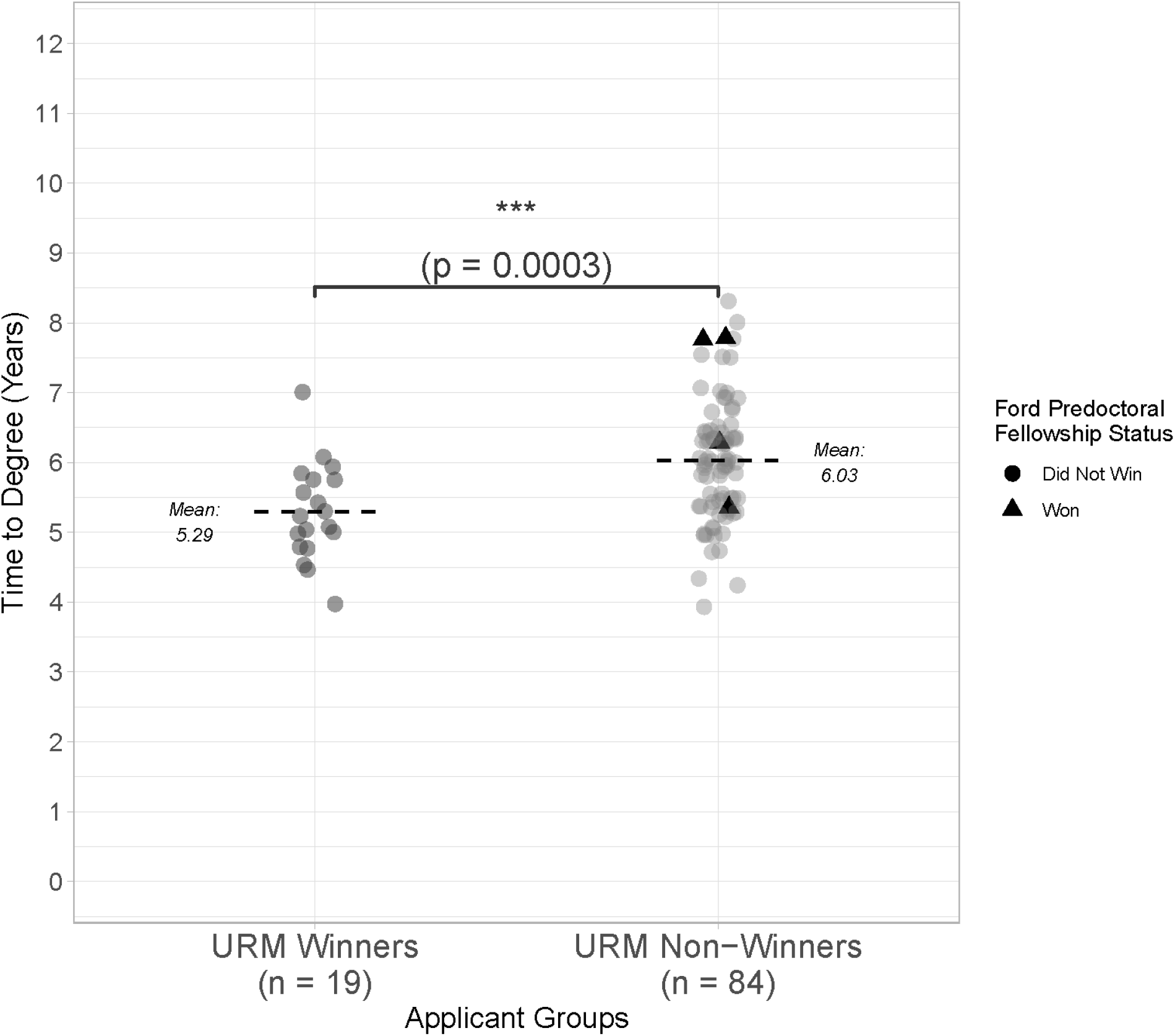
URM students who won grants (left) graduated faster than URM students who did not win grants (right). The Ford Fellowship winners (triangles) did not also win NSF or SeRA-tracked funding, were included within the URM non-winner group. The mean time to degree for each group is indicated by the dashed lines and the p-value is indicated by the asterisk at the top of the figure (p < 0.001).

The effect sizes of winning grants ranged from medium to large (using Cohen’s *d*). See Table 2 for a breakdown of the analyses and results with respect to winning grants. The largest effect was seen in the comparison between URM women who won awards and URM women who did not (1.25), once again highlighting that the relationship between time to degree and winning awards has a strong intersectional component, though in this case URM status (the second largest effect size, 0.88) is also an important dimension. Comparing URM and non-URM students yielded a medium effect size, suggesting that the stronger effects might be concentrated among just URM students.

**Table 2:** Effect sizes of winning a competitive award using Cohen’s *d* with corresponding figures and 95% confidence intervals.

| Comparison | Corresponding Figure | Cohen's $d$ | 95% Confidence Interval |
| --- | --- | --- | --- |
| URM winners vs. URM non-winners | 3 | 0.88 (large) | [0.36, 0.1.39] |
| URM winners vs. Non-URM winners | 2A | 0.54 (medium) | [0.02, 1.06] |
| URM Women winners vs. URM Women non-winners | 2B | 1.25 (large) | [0.46, 2.03] |

In the following section, we describe the results from our analysis of survey and interview data collected from current bioscience PhD students.

### 3.2 Qualitative results: Survey and interviews

We present the results for the qualitative analysis in two ways: proportions of responses to certain types of questions and with excerpts highlighting both broad themes and specific nuances in student experiences. Presenting the findings in this way will allow us to understand students’ decision-making processes with whether they apply for grants as well as ways they feel supported or not. “Support from advisors and mentors” was a key theme that emerged across the interviews, something we incorporate into the presentation of our results. Finally, because of identification risk and some of the interview participants representing very small populations, we limit the amount of detail we can share about the participants.

Among the 17 students interviewed, those who report having a more supportive advisor were more likely to apply for external funding (92%) than students who did not have supportive advisors (40%). See Figure 4 for a visualization of these disparities. All three URM women in the sample reported limits with the amount of support they received from advisors when it came to applying for awards; this was only the case for 14% of the rest of the sample who were not URM women. In fact, the only URM woman who applied and won an award participated in a supplemental mentorship program specifically designed to support URM students which she reported as an alternative source of support during her grant writing process. It should be noted that all the participants experienced the Covid-19 lockdowns early in their doctoral studies, yet 71% of the participants still reported having supportive advisors and the majority of the 29% who did not were URM women. Descriptively, these statistics indicate disparities in the quality of mentorship among bioscience doctoral students that are strongly correlated with whether or not they apply and eventually win competitive awards and funding. Given our longitudinal findings that URM women who secure awards graduate nearly a year faster than URM women who do not receive awards, it becomes something of concern if, as the interview data suggest, URM women are not receiving the same type of supportive mentorship that their peers receive which discourages them from applying.

**Fig. 4.**
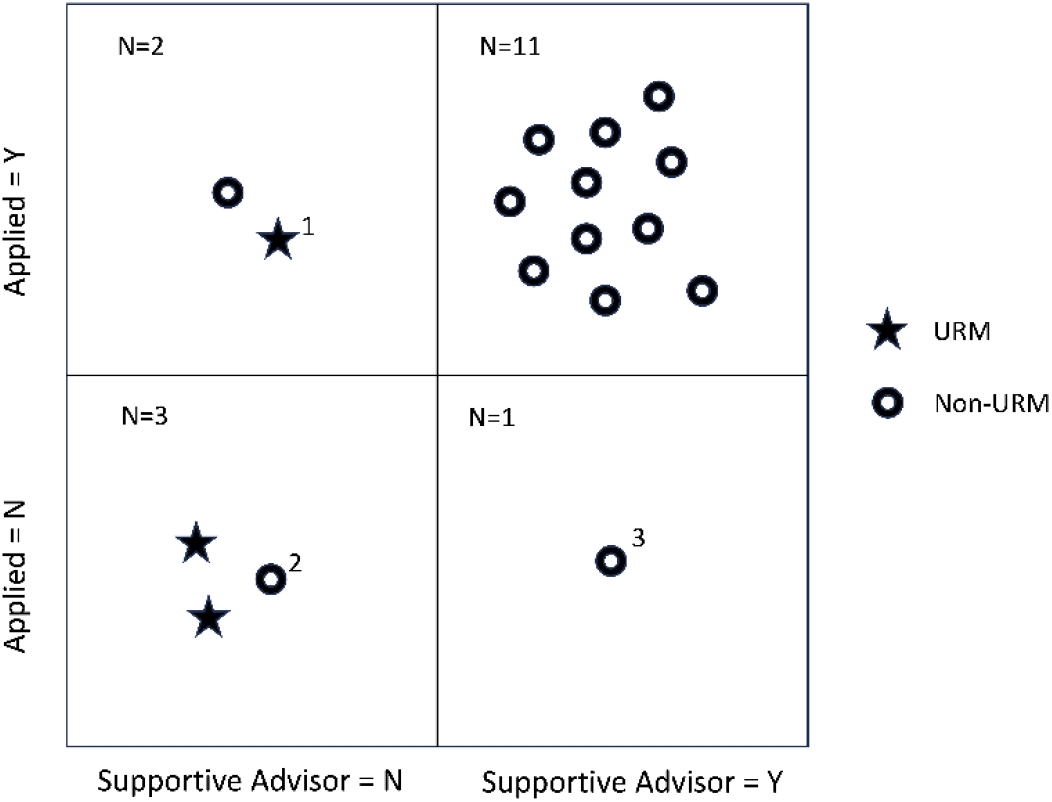
Students who did not have supportive mentors (left column) were less likely to apply for grants and other awards (top row). Students who identified as URM (stars) were less likely to have supportive advisors than other students (rings). 1: Had grant writing support outside of advisor. 2: Part of marginalized identity group not relating to race/ethnicity. 3: Student reported being unable to find relevant funding opportunities

In the interviews, the participants went into fine detail about how they understood and defined “support” in the context of grant writing as well as pointed to its crucial role in their academic successes. Table 2 presents examples of students describing the exact ways they were supported in grant writing and how they connect this support to the entire grant writing process. What became clear in the interviews is that students felt supported when advisors gave them direct, concrete feedback and attention.

For example, students describe “iterative” processes where they go back and forth with advisors at the “initial stages of brainstorming” all the way through going “back and forth” leading up to a submission. Even a student who ultimately did not win reported a similar intensive, iterative experience with their advisor. Collectively, these types of responses are examples of when doctoral students in the biosciences feel supported when their advisors are giving them sustained feedback and attention as they work through the grant writing process. In all of their answers, repetition and rewriting were also clear themes, pointing to another potentially positive way that advisors support their students with grant writing: encouraging revision and persistence. These students also had a higher likelihood of winning grants in the sample and were less likely to identify as URM. The clear examples and descriptions of support and their positive association with applying and winning awards contrast with students who felt less supported in their grant writing (see Table 3).

**Table 3:** Excerpts of students describing how they feel supported.

| Excerpt | Status |
| --- | --- |
| *We had like a brainstorming session, like pretty early on... he gave me a lot of feedback on... a method that I proposed, because obviously, I was new to the field at the time. So I didn't know like, what methods are feasible to be used to answer like a research question.* | Applied, received award. (non URM, Woman) |
| *And so he gave me a lot of ...constructive feedback on like, just methods that I could use or can think about to incorporate in this proposal idea. He also helped me draft sort of like, specific aims. So the way that I was writing my statement, before I got his help was like, not in a specific aims format. So he was able to help me... draft that sort of format, in a way that made it clear on what my research outline was. And then he kind of walked me through... how to formulate a feasible research question.* |  |
| *I had finished one version of a draft, I'd send it to her by email. And usually she just like, make comments on it and send it back. It is very iterative. But my advisor is generally very supportive, and very open. And I'd say like, most of the lab feels pretty comfortable just like talking about anything with her. So I'm talking about, you know, grant funding was, it was not a difficult thing to try to approach.* | Applied, received award. (non URM, Woman) |
| *And so they were helpful in the initial stages of brainstorming, you know, I had already kind of knew what my thesis project was going to be. So it was more a question of what is the best way to organize it into a format that fits with these grants, because different grants have different expectations. So that was the early stages of just like, what's the outline of it going to be? And then once I started writing it, then it was getting feedback from them... They're like, great.* | Applied, received award. (non URM, non Woman) |
| *Before sending in the application, I would send him my application for review. And I think like, for the first time that I was, I was submitting an application he sent me feedback, and then I improved it and send it back to him. And then he sent it back to me. So we're like, we went back and forth, like a couple of times. So it was really, really helpful.* | Applied, received award. (non URM, Woman) |
| *I would say like my PI was probably the primary source of mentorship. He's like, the person who like read everything over and like wrote the research statements and stuff with me.* | Applied, did not receive award. (non URM, Woman) |
| *...I would say he's fairly hands on... I know, some people kind of write it up and then the PI just kind of reads it over and then send you the edits. And then you're kind of like, that's kind of it. But like, we went through, like several rounds of writing together. And he wrote his own like, letter. He actually wrote it himself. And when I had trouble tracking down like faculty members for like, their letters, and supported my biosketch, he offered to help and kind of like to track them down as well. Yeah, so he was great.* |  |

**Table 4:** Excerpts of students describing how they do not feel supported.

| **Excerpt** | **Status** |
| --- | --- |
| *I am in a big lab, and my PI doesn't really often have time for sort of like hands on preparation for grants. And so I wanted that, and I wanted, you know, mechanism for feedback for my peers. And so I did that, to see if it would be something that I wanted to apply for.* | Applied, did not receive an award. (non URM, Woman) |
| *No one was like, you should apply to this. You should be doing this. Why don't you get this together? Like I feel like a lot of you don't like what constitutes good mentorship is telling people what they need to do that they don't need to know that they need to do and I got none... I think I was just very like, intimidated by the concept of applying for funding through an organization like the NIH. It's not like I had particular acute application* | Did not apply, did not receive an award. (non URM, Woman) |
| *trauma or anything like that. And certainly barely interacted with the NIH before... I think I just more psyched myself out of it more than anything else. And I don't think I'm alone in being intimidated by it. But I might be unusual in that I let it get the best of me...It's just so much stuff...I very much was guilty of not knowing what I didn't know. And not knowing that there were things that I didn't know that I needed to be on top of or find out... Like, there was no one to tell me all of this. My PI certainly didn't. And so I think if I had had that mentorship, I would have probably applied to all sorts of things. I just needed someone to tell me that I could.* |  |
| *And they are missing the fact that if I have to get a letter of rec from somebody who does not like me, for all these reasons that are masking their racist ideology... Like, how am I supposed to navigate applying for a grant when they are unconsciously going to sabotage what it is that I'm doing?* | Did not apply, did not receive an award.<br>(URM, Woman) |
| *I wouldn't say someone reached out or provided mentorship or anything like that. It was mainly resources and things that maybe I heard about or knew about, or other people had success about... Having other classmates ahead of me and telling me what works or not or me getting emails from, you know, the biosciences about different workshops and activities, and me actually taking the initiative to do those things myself. So I wouldn't say that someone encouraged me or told me to do anything. And as a woman of color... it is not the typical cadence of this community to always think of people of color first, or always include us on resources and things that could help them for this.* | Did not apply, did not receive an award.<br>(URM, Woman) |

Where the supported students describe the benefits of having an advisor who provides hands-on guidance, the students who did not feel supported noted how this was explicitly missing in their doctoral experiences. This includes instances where students reported being in a “big lab” and unable to find direct support from their PI to feeling actively discouraged due to their PI “not liking” them to the point of “unconsciously sabotaging” them. While we do not make any claims about the quality of mentorship nor the veracity of these claims, these quotes nevertheless highlight how feeling unsupported is a powerful influence of both perception and experience in doctoral education. Students also described not knowing where or how to even begin the process of writing a grant, “psyching” themselves out of beginning, and their PIs not alerting them to any relevant opportunities. Unsurprisingly, these students were also less likely to even apply for grants so much as win them. This, coupled with the fact that most of these students identified as either URM, women, or URM women points to a unique form of educational inequality to investigate further. Given that students identifying as URM women completed their PhD significantly faster when they also win research awards further justifies other institutions looking into their own data.

## 4 Discussion and Future Directions

We report on lessons learned from our longitudinal case study using 10 years of quantitative data and recent interview data. Using statistical (t-tests and Cohen’s *d*) and qualitative methods (thematic analysis), we found that faster times to degree are associated with winning competitive grants for specific groups of students, including URM students and URM women. Specifically, URM students who won research awards graduated faster than non-URM students who also won research awards. Likewise, we found that URM winners graduated faster than URM non-winners. When it comes to winning competitive research awards, the stakes are higher for URM women: when they won, they graduated almost a year faster (see Figure 2b). In the qualitative analysis, we found that currently enrolled students who identify as URM and/or women described feeling less supported in their grant writing. Conversely, students who described feelings of support were more likely to apply and win grants but also not identify as URM. Our case study points to important trends in graduate training at our study site (Stanford University) but also highlights the need for a cross-institutional study to see if these patterns are unique or pervasive.

While we do not make any causal claims about the relationship between winning competitive awards and time to degree, there are nevertheless important implications to consider for doctoral students in the biosciences. For URM and URM women students, the strength of the relationship between time to degree and awards as well as feelings of less advisor support necessitates further investigations to determine causality. It is possible that winning competitive awards serves as a strong validation of the research goals and interests, and scientific identity (Committee on Effective Mentoring in STEMM et al., 2019) for URM and URM women, which was shown to directly influence persistence in the sciences (Estrada et al., 2011). Furthermore, the hands-on support, such as iterative feedback during the grant writing process, may also validate scientific identify and shorten time to degree. However, the trend with the Ford winners shows that service and community engagement might delay times to degree. This delay is consistent with a common and well-studied phenomena often referred to as a cultural tax, minority tax, or invisible labor. This tax, while valuable to institutions, negatively impacts academic progress for individuals from marginalized identity groups (Cleveland et al., 2018; Padilla, 1994; Reid, 2021; Rodríguez et al., 2015). Likewise, other studies have suggested that racialized and gendered minorities seeking grants tend to propose research projects which directly address public health and community needs, topics which are less likely to get funding (Kolev et al., 2019). Future studies are needed to draw conclusive and generalizable results in order to tie such findings to ours.

Since many students are never formally trained in how to write proposals despite the practical benefits (Kulage et al., 2020; Mackert et al., 2017; von Hippel & von Hippel, 2015), these results point to practical ways to streamline the student experience by providing grant writing training (Botham et al., 2020). Doing so could have practical benefits, such as reducing time to degree. Because obtaining a first grant is associated with successfully acquiring future grants, an example of an academic Matthew effect (Bol et al., 2018), it would be interesting to investigate if grant writing successes as a graduate student correlate with future successes at the faculty stage. Future studies could take a deeper dive into these relationships and even analyze the types of grants that students are writing and submitting. Other follow-up studies may include tracking students who win competitive awards beyond their PhD to see if there is variation in career outcomes and filter the data and analysis by URM status and gender. The statistical and computational tools we utilized could be adapted for these analyses and help departments and institutions reconsider how they measure success and identify who benefits most from specific interventions.

The multiple methods approach which guided our analysis could also be used to compare the patterns in our dataset with other data across systems and organizations to improve educational outcomes and programming. These analyses could extend other educational practices and research experiences that have positive relationships between specific graduate experiences and career trajectories (Brazas & Ouellette, 2016). Beyond comparing data and patterns within them, future studies could also create interactive data dashboards to disseminate broadly. The potential use of these findings outside of purely academic research shows one way that researchers across titles and departments could participate in institutional analysis and improvement. For example, this work could become integral to other activities intended to improve educational experiences and outcomes for computational bioscientists (Atwood et al., 2015; Gurwitz et al., 2020). If the patterns we describe in our dataset mirror patterns at other schools, these findings could be important information that the entire discipline should consider and address (Blank et al., 2017). Such work would be of interest to other researchers but also to other educational stakeholders.

Considering the professional benefits of grant writing and our analysis showing its relationship with time to degree, we encourage URM and especially URM women to pursue competitive research awards. Of course, grant writing entails not just the act of writing a proposal but the entire process of designing a project and executing a research plan. Thus, grant writing is an important skill to develop earlier rather than later. Advisors should encourage their students to write grants as well as provide supportive, iterative feedback throughout that experience. In addition, it is important to ensure PhD advisors utilize effective research mentoring strategies, especially as their students are writing grants. For example, the Center for the Improvement of Mentored Experiences in Research has developed curricula specifically to accomplish this and has had some success (Handelsman et al., 2005). Advisors need to be informed about the impact grant writing (and potentially other skills often deemed as ‘peripheral’ for bioscience PhD students) may have on time to degree in order to provide advisee-specific advice that will advance their training and development into an independent and successful scientist.

## Declarations

### Data availability

An anonymized version of the data is available upon request to the corresponding authors.

### Ethical approval

The research done for this study followed the relevant guidelines and regulations to protect the anonymity of the students described in the data.

### Informed consent

We followed institutional IRB guidelines for our survey and interview study (IRB #65653).

## References

Akabas, M. H., & Brass, L. F. (2019). The national MD-PhD program outcomes study: Outcomes variation by sex, race, and ethnicity. JCI Insight, 4(19), e133010. https://doi.org/10.1172/jci.insight.133010

Allington, D., Hirsh, D., & Katz, L. (2023). Antisemitism is predicted by anti-hierarchical aggression, totalitarianism, and belief in malevolent global conspiracies. Humanities and Social Sciences Communications, 10(1), 155. https://doi.org/10.1057/s41599-023-01624-y

Atwood, T. K., Bongcam-Rudloff, E., Brazas, M. E., Corpas, M., Gaudet, P., Lewitter, F., Mulder, N., Palagi, P. M., Schneider, M. V., van Gelder, C. W. G., & GOBLET Consortium. (2015). GOBLET: The Global Organisation for Bioinformatics Learning, Education and Training. PLOS Computational Biology, 11(4), e1004143. https://doi.org/10.1371/journal.pcbi.1004143

Bernard, R. E., & Cooperdock, E. H. G. (2018). No progress on diversity in 40 years. Nature Geoscience, 11(5), 292–295. https://doi.org/10.1038/s41561-018-0116-6

Blank, R., Daniels, R. J., Gilliland, G., Gutmann, A., Hawgood, S., Hrabowski, F. A., Pollack, M. E., Price, V., Reif, L. R., & Schlissel, M. S. (2017). A new data effort to inform career choices in biomedicine. Science, 358(6369), 1388–1389. https://doi.org/10.1126/science.aar4638

Bloch, C., Graversen, E. K., & Pedersen, H. S. (2014). Competitive Research Grants and Their Impact on Career Performance. Minerva, 52(1), 77–96. https://doi.org/10.1007/s11024-014-9247-0

Bol, T., de Vaan, M., & van de Rijt, A. (2018). The Matthew effect in science funding. Proceedings of the National Academy of Sciences, 115(19), 4887–4890. https://doi.org/10.1073/pnas.1719557115

Botham, C. M., Brawn, S., Steele, L., Barrón, C. B., Kleppner, S. R., & Herschlag, D. (2020). Biosciences Proposal Bootcamp: Structured peer and faculty feedback improves trainees’ proposals and grantsmanship self-efficacy. PLOS ONE, 15(12), e0243973. https://doi.org/10.1371/journal.pone.0243973

Braun, V., & Clarke, V. (2012). Thematic analysis. In H. Cooper, P. M. Camic, D. L. Long, A. T. Panter, D. Rindskopf, & K. J. Sher (Eds.), APA handbook of research methods in psychology, Vol 2: Research designs: Quantitative, qualitative, neuropsychological, and biological. (pp. 57–71). American Psychological Association. https://doi.org/10.1037/13620-004

Brazas, M. D., & Ouellette, B. F. F. (2016). Continuing Education Workshops in Bioinformatics Positively Impact Research and Careers. PLOS Computational Biology, 12(6), e1004916. https://doi.org/10.1371/journal.pcbi.1004916

Cameron, C., Lee, H. Y., Anderson, C. B., Trachtenberg, J., & Chang, S. (2020). The role of scientific communication in predicting science identity and research career intention. PLOS ONE, 15(2), e0228197. https://doi.org/10.1371/journal.pone.0228197

Chaudhary, V. B., & Berhe, A. A. (2020). Ten simple rules for building an antiracist lab. PLOS Computational Biology, 16(10), e1008210. https://doi.org/10.1371/journal.pcbi.1008210

Cleveland, Dr. R., Sailes, Dr. J., Gilliam, Dr. E., & Watts, J. (2018). A Theoretical Focus on Cultural Taxation: Who Pays for It in Higher Education. Advances in Social Sciences Research Journal, 5(10). https://doi.org/10.14738/assrj.510.5293

Cohen, J. (2013). Statistical power analysis for the behavioral sciences. Academic press. Academic Press.

Committee on Effective Mentoring in STEMM, Board on Higher Education and Workforce, Policy and Global Affairs, & National Academies of Sciences, Engineering, and Medicine. (2019). The Science of Effective Mentorship in STEMM (A. Byars-Winston & M. L. Dahlberg, Eds.; p. 25568). National Academies Press. https://doi.org/10.17226/25568

de Valero, Y. F. (2001). Departmental Factors Affecting Time-to-Degree and Completion Rates of Doctoral Students at One Land-Grant Research Institution. The Journal of Higher Education, 72(3), 341–367. https://doi.org/10.1080/00221546.2001.11777098

Ehrenberg, R. G., & Mavros, P. G. (1995). Do Doctoral Students’ Financial Support Patterns Affect Their Times-To-Degree and Completion Probabilities? The Journal of Human Resources, 30(3), 581. https://doi.org/10.2307/146036

Estrada, M., Woodcock, A., Hernandez, P. R., & Schultz, P. W. (2011). Toward a model of social influence that explains minority student integration into the scientific community. Journal of Educational Psychology, 103(1), 206–222. https://doi.org/10.1037/a0020743

Feldon, D. F., Peugh, J., Maher, M. A., Roksa, J., & Tofel-Grehl, C. (2017). Time-to-Credit Gender Inequities of First-Year PhD Students in the Biological Sciences. CBE—Life Sciences Education, 16(1), ar4. https://doi.org/10.1187/cbe.16-08-0237

Files, D. C., Hume, P. S., Krall, J., Montemayor, K., Schmidt, E. P., & King, L. S. (2020). Grant Writing for Clinicians in Training. Chest, 157(4), 932–935. https://doi.org/10.1016/j.chest.2019.10.024

Gurwitz, K. T., Singh Gaur, P., Bellis, L. J., Larcombe, L., Alloza, E., Balint, B. L., Botzki, A., Dimec, J., Dominguez del Angel, V., Fernandes, P. L., Korpelainen, E., Krause, R., Kuzak, M., Le Pera, L., Leskošek, B., Lindvall, J. M., Marek, D., Martinez, P. A., Muyldermans, T., … Rustici, G. (2020). A framework to assess the quality and impact of bioinformatics training across ELIXIR. PLOS Computational Biology, 16(7), e1007976. https://doi.org/10.1371/journal.pcbi.1007976

Handelsman, J., Handelsman, H., Wisconsin Program for Scientific Teaching, & Howard Hughes Medical Institute Professors Program (Eds.). (2005). Entering mentoring: A seminar to train a new generation of scientists. Board of Regents of the University of Wisconsin System.

Hofstra, B., Kulkarni, V. V., Munoz-Najar Galvez, S., He, B., Jurafsky, D., & McFarland, D. A. (2020). The Diversity–Innovation Paradox in Science. Proceedings of the National Academy of Sciences, 117(17), 9284–9291. https://doi.org/10.1073/pnas.1915378117

Kahn, R., Conn, G., Pavlath, G., & Corbett, A. H. (2016). Use of a grant writing class in training PhD students. 17(7). https://doi.org/10.1111/tra.12398

Kolev, J., Fuentes-Medel, Y., & Murray, F. (2019). Is Blinded Review Enough? How Gendered Outcomes Arise Even Under Anonymous Evaluation (No. w25759; p. w25759). National Bureau of Economic Research. https://doi.org/10.3386/w25759

Kulage, K. M., Stone, P. W., & Smaldone, A. M. (2020). Supporting dissertation work through a nursing PhD program federal grant writing workshop. Journal of Professional Nursing, 36(2), 29–38. https://doi.org/10.1016/j.profnurs.2019.08.001

Kumari, J., & Kumar, J. (2023). Influence of motivation on teachers’ job performance. Humanities and Social Sciences Communications, 10(1), 158. https://doi.org/10.1057/s41599-023-01662-6

Mackert, M., Donovan, E. E., & Bernhardt, J. M. (2017). Applied Grant Writing Training for Future Health Communication Researchers: The Health Communication Scholars Program. Health Communication, 32(2), 247–252. https://doi.org/10.1080/10410236.2015.1110686

Master, A., & Meltzoff, A. N. (2020). Cultural Stereotypes and Sense of Belonging Contribute to Gender Gaps in STEM. International Journal of Gender, Science and Technology, 12(1), 152–198.

McCoy, D. L., Luedke, C. L., & Winkle-Wagner, R. (2017). Encouraged or Weeded Out: Perspectives of Students of Color in the STEM Disciplines on Faculty Interactions. Journal of College Student Development, 58(5), 657–673. https://doi.org/10.1353/csd.2017.0052

Mendoza-Sanchez, I., deGruyter, J. N., Savage, N. T., & Polymenis, M. (2022). Undergraduate GPA Predicts Biochemistry PhD Completion and Is Associated with Time to Degree. CBE—Life Sciences Education, 21(2), ar19. https://doi.org/10.1187/cbe.21-07-0189

Ostriker, J., Holland, P., Kuh, C. V., & Voytuk, J. A. (2010). A Data-Based Assessment of Research-Doctorate Programs in the United States. http://www.nap.edu/catalog/12850.html

Padilla, A. M. (1994). Research news and Comment: Ethnic Minority Scholars; Research, and Mentoring: Current and Future Issues. Educational Researcher, 23(4), 24–27. https://doi.org/10.3102/0013189X023004024

Palmer, R., Maramba, D., & Dancy II, T. (2011). A Qualitative Investigation of Factors Promoting the Retention and Persistence of Students of Color in STEM. Journal of Negro Education, 80(4).

Petersen, S. L., Erenrich, E. S., Levine, D. L., Vigoreaux, J., & Gile, K. (2018). Multi-institutional study of GRE scores as predictors of STEM PhD degree completion: GRE gets a low mark. PLOS ONE, 13(10), e0206570. https://doi.org/10.1371/journal.pone.0206570

PREDOCTORAL FACT SHEET. (2023). The National Academies of Sciences, Engineering, and Medicine. https://sites.nationalacademies.org/PGA/FordFellowships/PGA_047958

Price, J. (2006). Does a Spouse Slow You Down?: Marriage and Graduate Student Outcomes. SSRN Electronic Journal. https://doi.org/10.2139/ssrn.933674

Quitadamo, I. J., & Kurtz, M. J. (2007). Learning to Improve: Using Writing to Increase Critical Thinking Performance in General Education Biology. CBE—Life Sciences Education, 6(2), 140–154. https://doi.org/10.1187/cbe.06-11-0203

Ransdell, L., Lane, T., Schwartz, A., Wayment, H., & Baldwin, J. (2021). Mentoring New and Early-Stage Investigators and Underrepresented Minority Faculty for Research Success in Health-Related Fields: An Integrative Literature Review (2010–2020). International Journal of Environmental Research and Public Health, 18(2), 432. https://doi.org/10.3390/ijerph18020432

Reid, R. A. (2021). Retaining Women Faculty: The Problem of Invisible Labor. PS: Political Science & Politics, 54(3), 504–506. https://doi.org/10.1017/S1049096521000056

Rodríguez, J. E., Campbell, K. M., & Pololi, L. H. (2015). Addressing disparities in academic medicine: What of the minority tax? BMC Medical Education, 15(1), 6. https://doi.org/10.1186/s12909-015-0290-9

Sargent, J., Williams, R. A., Hagerty, B., Lynch-Sauer, J., & Hoyle, K. (2002). Sense of Belonging as a Buffer Against Depressive Symptoms. Journal of the American Psychiatric Nurses Association, 8(4), 120–129. https://doi.org/10.1067/mpn.2002.127290

Smith, J. L., Stoop, C., Young, M., Belou, R., & Held, S. (2017). Grant-Writing Bootcamp: An Intervention to Enhance the Research Capacity of Academic Women in STEM. BioScience, 67(7), 638–645. https://doi.org/10.1093/biosci/bix050

Strayhorn TL. (2018). College students’ sense of belonging: A key to educational success for all students.

Sullivan, G. M., & Feinn, R. (2012). Using Effect Size—Or Why the P Value Is Not Enough. Journal of Graduate Medical Education, 4(3), 279–282. https://doi.org/10.4300/JGME-D-12-00156.1

Thorpe, R. J., Vishwanatha, J. K., Harwood, E. M., Krug, E. L., Unold, T., Eide Boman, K., & Jones, H. P. (2020). The Impact of Grantsmanship Self-Efficacy on Early Stage Investigators of The National Research Mentoring Network Steps Toward Academic Research (NRMN STAR). Ethnicity & Disease, 30(1), 75–82. https://doi.org/10.18865/ed.30.1.75

von Hippel, T., & von Hippel, C. (2015). To Apply or Not to Apply: A Survey Analysis of Grant Writing Costs and Benefits. PLOS ONE, 10(3), e0118494. https://doi.org/10.1371/journal.pone.0118494

Walton, G. M., & Cohen, G. L. (2007). A question of belonging: Race, social fit, and achievement. Journal of Personality and Social Psychology, 92(1), 82–96. https://doi.org/10.1037/0022-3514.92.1.82

Walton, G. M., & Cohen, G. L. (2011). A Brief Social-Belonging Intervention Improves Academic and Health Outcomes of Minority Students. Science, 331(6023), 1447–1451. https://doi.org/10.1126/science.1198364

Walton, G. M., Cohen, G. L., Cwir, D., & Spencer, S. J. (2012). Mere belonging: The power of social connections. Journal of Personality and Social Psychology, 102(3), 513–532. https://doi.org/10.1037/a0025731

